# β_2_-Adrenergic Signaling Switches from Cardioprotective to Cardiotoxic in Acute vs. Chronic Oxidative Stress

**DOI:** 10.64898/2026.04.22.720269

**Authors:** Giovanni Fajardo, Mingming Zhao, Gwanghyun Jung, Viswanathan Rajagopalan, Michael Coronado, Sushma Reddy, Daniel Bernstein

## Abstract

**BACKGROUND AND PURPOSE:** β-adrenergic receptors (AR) regulate both cardiac function and remodeling. Many studies suggest that, in addition to their effects on heart rate and contractility, β_1_-ARs mediate cardiotoxic signaling, whereas β_2_-ARs are generally cardioprotective. However, there is conflicting data on the role of β_2_-ARs, differing dependent on the nature of the stress. Given the extremely common use of β-blockers and agonists clinically, we sought to understand the differential cardioprotective/cardiotoxic effects of β_2_-AR signaling dependent on timing (acute vs. chronic) and type of cardiotoxic stress.

**EXPERIMENTAL APPROACH:** Wild-type (WT) and β-AR knockout (β_1_-KO and β_2_-KO) mice were subjected to acute (15 mg·kg^-1^ x 1 dose) or chronic (2 mg·kg^-1^·wk^-1^ x 7 wks) oxidative stress using doxorubicin (DOX). Survival, cardiac function and histopathology were assessed and differential signaling activation determined by Western blot and gene expression by RNA-seq.

**KEY RESULTS:** We have shown that β_2_-KOs manifest extreme cardiotoxicity with acute DOX (100% mortality within 30 min), supporting a strong cardioprotective role of β_2_-signaling. In marked contrast, with chronic DOX, β_2_-KO had enhanced survival (t½ 54 d vs. 42 d in WT) and attenuated cardiac dysfunction. In β_2_-KO, acute DOX activated stress MAPKs (p38, ERK and JNK), whereas chronic DOX did not; furthermore, in the absence of β2-ARs, oxidative stress and lipid accumulation were reduced, genes regulating compensatory metabolic pathways (AMPK and insulin/PI3K) were upregulated, and genes regulating mitochondrial and contractile function were preserved, whereas they were downregulated in WT with chronic DOX.

**CONCLUSIONS:** β_2_-AR signaling switches from being cardioprotective during acute oxidative stress, to cardiotoxic during chronic stress. Inhibition of β_2_-AR signaling during chronic stress induces signaling and metabolic compensations that serve to reduce oxidative injury. This unexpected temporal switching has potential significant implications for all models of cardiovascular disease, as well as for the clinical use of subtype-specific β-blockers.

**CLINICAL PERSPECTIVE:** *What is new?:*

- Our finding that β2-adrenergic receptor signaling can switch from being beneficial (cardioprotective) to detrimental (cardiotoxic) depending on the acuteness or chronicity of a cardiac stressor.
- Identification of the mechanisms by which this temporal switch is mediated could lead to new drug development.

*What are the clinical implications?:*

- Our findings provide potential guidance in choosing between a β1-specific vs. a β1/2-non-specific drug when treating specific cardiovascular diseases based on their temporal characteristics.
- The temporal protective/toxic switching that we describe could be a mechanism common to many other drugs, yet is rarely tested, suggesting the need for additional studies using temporal course as a factor.

## INTRODUCTION

β-adrenergic receptor (β-AR) signaling is one of the primary regulators of cardiac function, but also mediates adverse remodeling in heart failure associated with sustained sympathetic activation (1). β-blockers are thus one of the top ten prescribed drugs both in the US and globally, and β-agonists one of the most utilized agents in patients with pulmonary disease and in those in the intensive care unit. β_1_-ARs are subtype-selectively downregulated in heart failure, and β-antagonist therapy (either β_1_-specific blockers such as metoprolol or β_1_β_2_ blockers such as carvedilol) is now standard therapy to reduce sympathetic-induced injury and restore normal β-AR signaling homeostasis (2, 3). Accumulating data suggest that, in addition to its effects on inotropy and chronotropy, the β_1_-AR also induces cardiotoxic signaling, whereas the β_2_-AR induces cardioprotective signaling, as well as opposing β_1_-mediated cardiotoxic signaling through activation of Gi and by crosstalk with CaMKII, MAPKs and PI3K-Akt (4–7).

We have previously described a compelling differential role for β_1_ vs. β_2_-AR subtypes in mediating the effects of the anthracycline anti-tumor drug doxorubicin (DOX). Wildtype (WT) or β_1_-AR knockout (β_1_-KO) mice receiving a single, acute, high dose of DOX (15 mg·kg^-1^) do not manifest immediate cardiotoxicity but gradually develop dilated cardiomyopathy over the following 2-6 weeks. In contrast, β_2_-AR knockout (β_2_-KO) mice demonstrate markedly enhanced cardiotoxicity, leading to 100% mortality within 30 min, a dramatic response which we have also recapitulated in cardiomyocytes cultured from these mice (6, 8–10). Others have shown that mice overexpressing cardiac β_2_-ARs have enhanced preservation of left ventricular (LV) function following myocardial infarction (MI)(11), and that combination therapy with a β_1_-antagonist and a β_2_-agonist had a synergistic effect on survival after MI, reducing scar expansion and improving LV function (12). In combination, these studies provide additional support for a beneficial role of β_2_-signaling in several different models.

However, there is also conflicting data that suggests that β_2_-AR signaling may not always be cardioprotective and may even be cardiotoxic. At baseline, β_2_-KO mice have no significant alterations in cardiovascular function (13). Yet we have shown that deletion of the β_2_-AR rescues the cardiomyopathy induced by knockout of Muscle LIM Protein (MLP), mainly by improving Ca^2+^ transients. Knockout of the β_2_-AR has a similarly beneficial effect in pressure overload-induced heart failure induced by transverse aortic constriction (TAC) (14). In the same manner, cardiomyocyte specific overexpression of β_2_-ARs exaggerates TAC induced hypertrophy and worsens heart function (15, 16). Finally, in a model of diabetic cardiomyopathy, deletion of β_2_-signaling prevents myocardial injury and dysfunction (17). These data suggest that β_2_-ARs can also play a deleterious role during some types of cardiovascular stress.

The purpose of the current study was to understand the complex dual nature of β_2_-AR signaling in the context of acute vs. chronic cardiac stress, and to examine downstream signaling pathways to determine the mechanisms responsible for temporal differences in β-AR signaling. We demonstrate a unique switch in β_2_-AR signaling from *cardiotoxic* during acute DOX to *cardioprotective* during chronic DOX treatment. We conclude that the β_2_-AR’s role in cardiac remodeling is more complex than previously appreciated and highly context dependent: not only on the type of stress (ischemia/reperfusion, pressure overload, genetic cardiomyopathy, oxidative cardiotoxic drug) but on the temporal nature of that stress (acute vs. chronic). A better understanding of how β_2_-AR signaling is regulated during these different conditions will allow the better application of β_2_-agonist or antagonist therapies tailored to each patient’s pathophysiology.

## METHODS

### Animals

Congenic FVB β_1_-KO and β_2_-KO mice were generated as previously described (18, 19). DOX was administered to β_1_-KO and β_2_-KO mice (3 months old, male) and wildtype (WT) age and strain-matched controls using either acute or chronic protocols. All procedures were performed in accordance with National Institutes of Health standards and were approved by the Administrative Panel on Laboratory Animal Care at Stanford University.

### Model of toxic cardiomyopathy

We used a murine model of chronic doxorubicin cardiotoxicity, based on established protocols (20, 21). Mice were treated with 2 mg·kg^-1^·wk^-1^ i.p. doxorubicin (NovaPlus, Bedford, OH) for 7 wks. For acute doxorubicin cardiotoxicity, we injected a single dose of 15 mg·kg^-1^ of doxorubicin (200–300 microliters) via the dorsal tail vein (8). The doses we used for both acute (22–25) and chronic (20, 21, 26–29) studies are based on an extensive literature in the mouse, both of which have been shown to induce a dilated cardiomyopathy with different time courses. We also verified the appropriateness of these dosing regimens to match clinical oncology regimens using two methods. First, comparing serum levels, doses in humans range from 20-30 mg·(m^2^)^-1^ given weekly to 35-90 mg·(m^2^)^-1^ given every 3-4 weeks, yielding levels at 30 min of 0.6-2.5 µg· ml^-1^. In mice given 6 mg·kg^-1^ i.v., levels at 30 min were 0.6 µg·ml^-1^ (30, 31). Mice given 15 mg·kg^-1^ i.v. had levels at 30 min of 1.0-2.0 µg·ml^-1^ (30, 32). Second, we calculated the human equivalent dose (HED) based on either body weight or surface area. FDA guidelines for conversion of murine doses to HED are based on BSA (Center for Drug Evaluation and Research) for which a murine dose of 8.5 mg·kg^-1^ is equivalent to 25 mg·(m^2^)^-1^ in humans (33). Thus, the low, chronic dose we used in mice should result in a serum level about half of that obtained with the lower-end therapeutic dose in humans (20 mg·(m^2^)^-1^) and, based on the murine literature, lead to the gradual development of a dilated cardiomyopathy. We also performed studies showing no difference based on whether doxorubicin was given i.v. or i.p. Finally, to verify that the intraperitoneal injections were not leading to peritonitis, serial post-mortem studies excluded both chemical and infectious peritonitis. Serial white blood counts (WBCs) were also performed to evaluate immune status, and these showed no significant changes. Thus, there was no confounding life-threatening pathology in these mice other than that related to their cardiomyopathy.

### Echocardiographic evaluation

Mice were studied at baseline and every 2 weeks under light anesthesia with isoflurane (induction 3%, maintenance 1.5%). 2D images in the parasternal short axis were obtained with a GE Vivid 7 ultrasound system (GE Health Care, Milwaukee, WI) equipped with a 13 MHz transducer. Left ventricular end-systolic (LVESD) and end-diastolic (LVEDD) dimensions were measured and left ventricular fractional shortening (FS) was calculated. Importantly, the echocardiographer was blinded to the experimental group.

### Protein expression and phosphorylation

Left ventricular (LV) myocardium was obtained at 20 min after DOX in the acute study and after 7 wks in the chronic study. Protein expression was determined by Western blot. Tissues were harvested and homogenized using lysis buffer (NaCl 80 mM·L^-1^, Triton X 0.05%, EDTA 1 mM/L, HEPES 20 mM·L^-1^, Sodium deoxycholate 0.5% deoxycholic acid, sodium salt stored at 4°C, including DTT 1 mM/L, β–glycerophosphate 20 mM·L^-1^ with TissueLyser LT (Qiagen Inc., Valencia, CA) and centrifuged at 10,000 rpm for 15 min at 4°C. Proteins were quantified with Bradford (Bio-Rad, Hercules, CA, USA) and separated by TrisGlycine SDS/PAGE (Bio-Rad). Transferred proteins on nitrocellulose membrane were probed against phosphorylated ERK1/2, p38 MAPK, JNK, and MKK4, obtained from Cell Signaling Technologies Inc. (Danvers, MA). Specific antibody against CaMKIIδ was gifted by Dr. Don Bers (UC Davis) and antibodies for total CaMKII and Bcl2 were from Santa Cruz Biotechnology Inc. (Santa Cruz, CA), phospho-CaMKII (T286, MA1-047) from Thermo Fisher Scientific Inc. (Rockford, IL). Phospho-AMPK (T172), and total-AMPK antibodies from Cell Signaling Technologies Inc. (Danvers, MA). HNE antibody was obtained from Alpha Diagnostics (San Antonio, TX).

### Electron microscopy

LV myocardium was excised and mitochondrial and sarcomeric morphology was assessed by transmission electron microscopy (TEM) as described previously (34). Lipid droplets were quantified and mitochondrial number and morphology (number, area, and perimeter) determined using Image J. From each tissue section, 10 fields were captured, measured and averaged for each mouse. Statistical comparisons were then performed between mice, resulting in a total n of 3-5 in each experimental group.

### RNA-seq analysis

RNA-seq analysis was performed on whole hearts from four WT and four β_2_-KO mice at baseline. Because β_2_-KO mice died within 15-30 min after acute high dose DOX, too soon to capture significant transcriptomic changes, we used an intermediate acute dose of 4 mg·kg^-1^ for 2 d, which we will refer to as “subacute” to examine mRNA changes after a large but sub-lethal acute dose. Three DOX-treated β_2_-KO were studied using this subacute protocol and three using the chronic protocol (2 mg·kg^-1^·wk^-1^ for 7 wks) (Figure 1B). Hearts were flash-frozen in liquid nitrogen and stored at −80 °C. RNA was extracted using a micro RNeasy kit (QIAGEN, Venlo, Netherlands), library preparation and sequencing on a HiSeq 4000 instrument (Illumina) was performed by Macrogen, Inc. (Seoul, South Korea). 40 million reads (20 million in each direction) were used for each sample. Reads were mapped to the mm10 reference mouse genome using TopHat2 software, assembled into transcripts using HTSeq-count and quantified using DESeq2. Only transcripts mapped to unique genes were retained and only known protein-coding genes were retained for subsequent analysis. Functional enrichment analyses of multiple group genes and genes-concept network analyses were performed using Ingenuity Pathway Analysis (IPA, www.qiagen.com/ingenuity, QIAGEN). The particularly grouped gene list was extracted from IPA.

**Figure 1:**
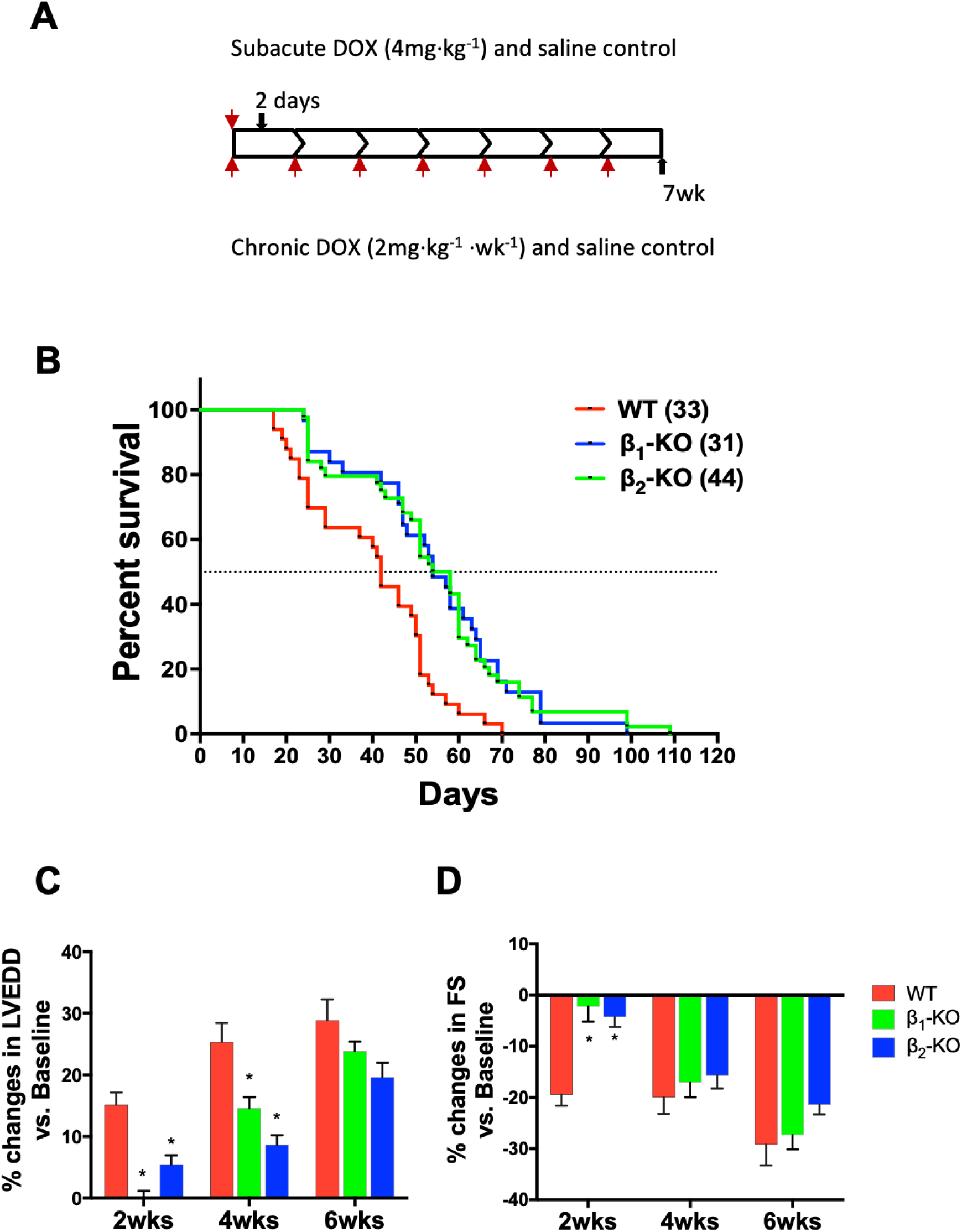
Beneficial effect of β_1_-and β_2_-adrenergic receptor knockout on survival and cardiac function with chronic DOX treatment. **A**. Schematic plan for RNA-seq analysis. DOX was injected either subacute (4mg·kg-1 for 2 days) or chronic (2mg·kg-1·wk-1 for 7wks), in WT and β2-KO mice. **B.** Enhanced survival in β_1_-KO and β_2_-KO mice after chronic DOX. WT, β_1_-KO and β_2_-KO mice were injected with chronic low-dose DOX and monitored for survival. WT had a median survival of 42d with 100% mortality by 70d. In contrast, survival was significantly enhanced in β_1_-KO (median survival 52d, 100% mortality at 99d) and in β_2_-KO (median survival 58d, 100% mortality at 109d). Number of mice shown in parentheses; Log rank P<0.0001. **C** and **D**. Slower progression of cardiac dysfunction in β-KO mice. Transthoracic echocardiography was performed serially in WT and β-KO mice following initiation of DOX treatment. (**C**) Slower progression of increase in LVEDD in both β-KOs; (**D**) Slower progression of decrease in fractional shortening (FS) in both β-KOs. Data show mean ± SEM, *p < 0.05 vs. corresponding weeks in WT group (two-way ANOVA with Tukey test).

### Data resources

## The accession number for the RNA-seq data reported in this paper is GEO

### Statistical analysis

All values shown are mean ± SEM. Statistical significance of differences in parameters was determined using Student’s T-test or for multiple comparisons with one way or two-way ANOVA with Bonferroni or Dunnett’s post-hoc tests where appropriate using GraphPad Prism (GraphPad Software Inc., La Jolla, CA). Actuarial survival was assessed by the Kaplan-Meier method using log rank testing. p<0.05 was considered significant

## RESULTS

### Enhanced survival and slower progression of LV dysfunction in both β_1_ and β_2_-KO mice following chronic DOX treatment

WT mice treated with chronic DOX developed dilated cardiomyopathy. Mortality in WT mice began at 33 d following initiation of treatment, with a median survival of 42 d and 100% mortality by 70 d. In contrast, deletion of either the β_1_-AR or the β_2_-AR resulted in significantly enhanced survival following chronic DOX treatment compared to WT (log rank p<0.0001). The median survival was enhanced in β_1_-KO to 54 d with 100% mortality by 99 d and the median survival in β_2_-KO was enhanced to 56 d with 100% mortality by 109 d (Figure 1B).

Serial echocardiograms, performed at 2-week intervals, showed left ventricle (LV) dilation, with increased LVEDD by 2 and 4 weeks in WT. In contrast, there was significantly attenuated dilation and delay in both knockouts (WT: 15.2±1.9% and 25.4±2.9%, β_1_-KO: 0.2±1.0% and 14.6±1.8%, β_2_-KO: 5.4±1.5% and 8.7±1.6%) (Figure 1B). Correspondingly, WT mice demonstrated a significant decrease in left ventricular fractional shortening (FS) by 2 weeks. In contrast, neither β_1_-KO nor β_2_-KO showed a significant change in FS at 2 weeks (WT:-19.5±2.2%, β_1_-KO:-2.2±2.9%, β_2_-KO: - 4.3±1.9%). Subsequently, both LVEDD and FS deteriorated further in the WT mice. In contrast, both β-AR knockouts demonstrated a significantly slower progression of cardiac dysfunction (vs. respective WT groups and earlier time points). Although the β_2_-KO showed a trend toward a slower progression than the β_1_-KO, the differences between the two knockouts were not statistically significant (Figure 1C-D). Lung weight to body weight ratio, a surrogate of heart failure, was increased by over 19% (p<0.05) at 7 weeks following DOX in both WT and β_1_-KO mice compared to baseline. In contrast, this ratio was not significantly altered in β_2_-KO mice at 7 weeks (Supporting Information Figure S1). These results indicate that deletion of either β-AR subtype confers a modest degree of cardioprotection during chronic low-dose treatment with DOX. Taken together with our previous data during acute DOX (6, 8–10), these results demonstrate that β_1_-ARs play a cardiotoxic role in both acute and chronic DOX-induced cardiotoxicity, however, β_2_-ARs play dramatically different roles in DOX-induced cardiotoxicity depending on the chronicity (Table 1), with the β_2_-KO driving 100% early lethality in acute DOX, converting to a moderate rescue effect in chronic DOX.

**Table 1.**
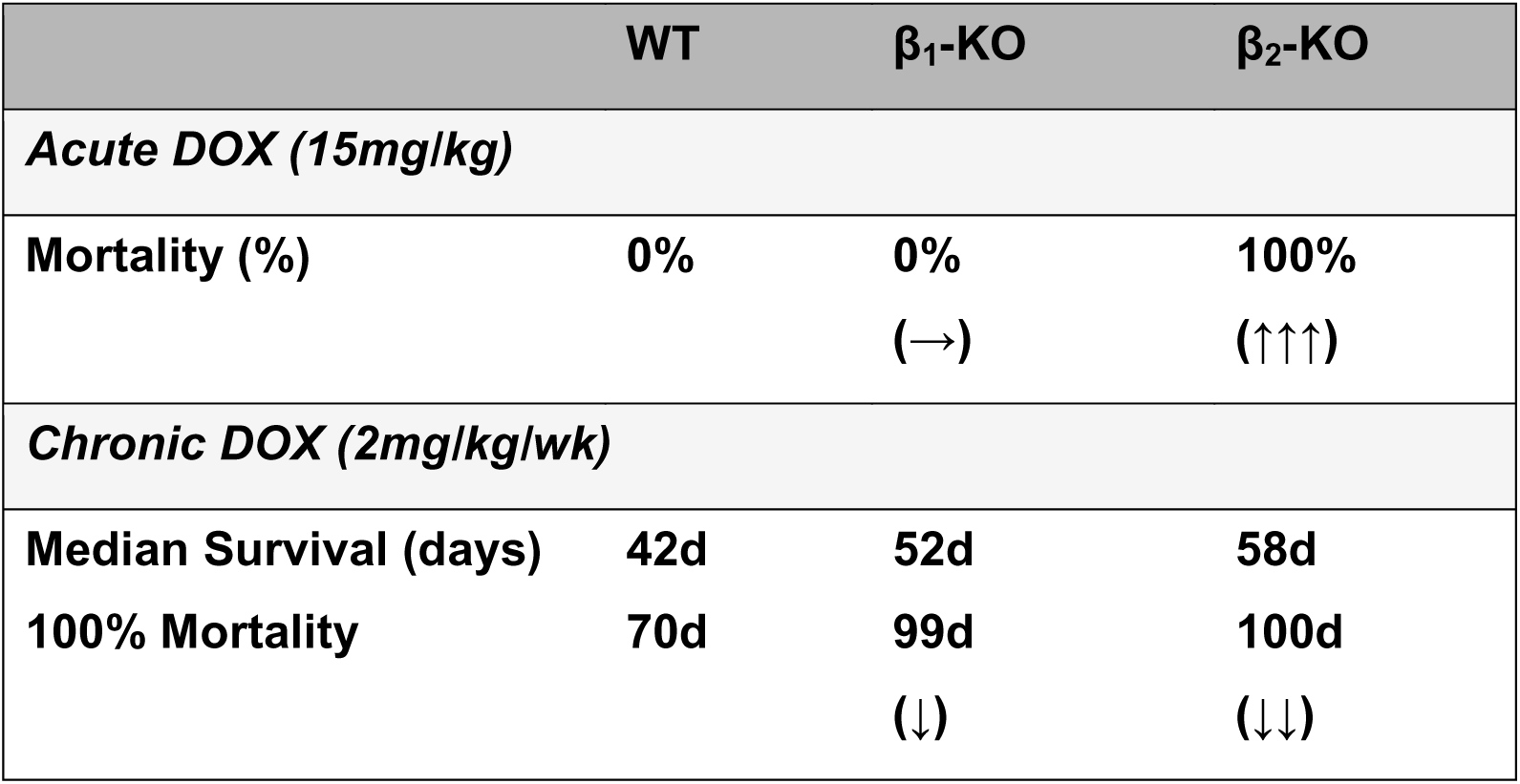
Comparison of temporal effects (acute vs chronic) of DOX administration on mortality in β_1_-KO and β_2_-KO mice. The β_2_-AR is the predominant temporal-switching regulator for flipping the response to acute DOX vs. Chronic DOX. Arrows show the direction of changes compared to WT and the number of arrows represents the magnitude of change.

| | WT | $\beta_1$-KO | $\beta_2$-KO |
| --- | --- | --- | --- |
| ***Acute DOX (15mg/kg)*** |  |  |  |
| **Mortality (%)** | **0%** | **0%**<br>(→) | **100%**<br>(↑↑↑) |
| ***Chronic DOX (2mg/kg/wk)*** |  |  |  |
| **Median Survival (days)** | **42d** | **52d** | **58d** |
| **100% Mortality** | **70d** | **99d**<br>(↓) | **100d**<br>(↓↓) |

### Stress MAPK activation with acute but not with chronic DOX in β_2_-KO mice

The opposing roles of the β_2_-AR in acute vs. chronic DOX stress prompted us to investigate the molecular mechanisms behind this dramatic switch. We have previously shown that three members of the MAPK family are highly activated following acute DOX treatment (15 mg·kg^-1^) in β_2_-KO mice compared to WT: p38, ERK1/2, and JNK, with p38 the most differentially upregulated (8). In contrast, with chronic DOX, β_2_-KO mice did not show activation of any MAPK family member (p38, ERK1/2, JNK) at 7 weeks following low-dose DOX treatment (Figure 2). Although MKK4, the kinase that phosphorylates JNK, was activated in the acute DOX treated groups, it was not activated following chronic DOX treatment.

**Figure 2.**
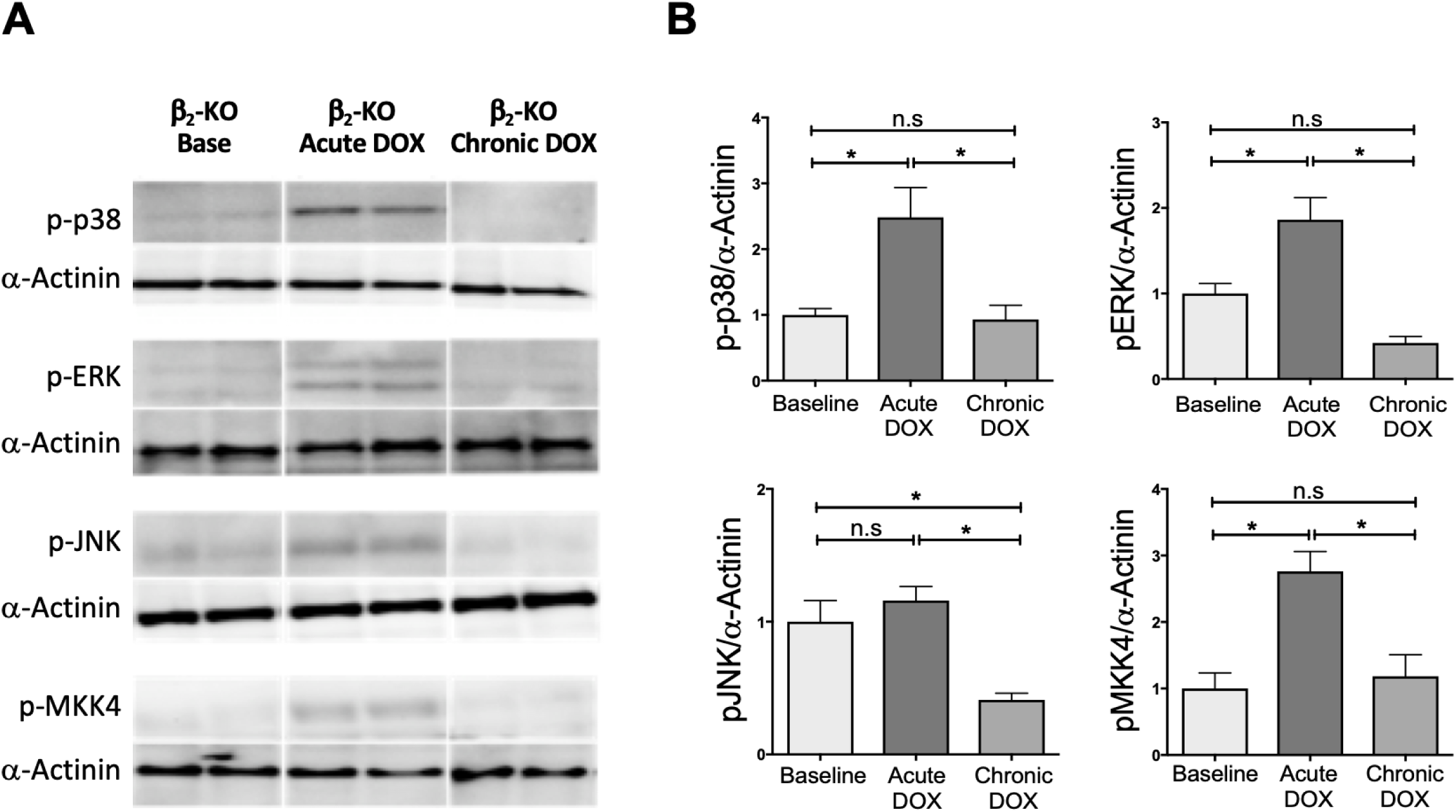
Stress response MAPK signaling is activated in acute but not in chronic DOX-treated β_2_-KO. Phosphorylation profiles of MAP kinases (p-38, p-ERK, and p-JNK) and MAP kinase kinase (MKK4) were detected by western blot. Representative blots (**A**) and quantified phosphorylation normalized by 𝛼-actinin are shown in **B**. Data show mean ± SEM, n≧4 per group, * *p*<0.05 and n.s, not significant (one-way ANOVA with Tukey test).

### Attenuated mitochondrial injury and lipid peroxidation with chronic DOX in β_2_-KO mice

Mitochondrial injury is one of the major mechanisms of DOX cardiotoxicity. We therefore examined mitochondrial structural integrity by electron microscopic (EM) analysis of LV myocardium. After chronic DOX treatment, mitochondrial disarray was seen in both WT and β_2_-KO compared to non-treated controls, which showed more regular alignment of mitochondria with sarcomere structure (Figure 3A) as previously described (35, 36). As has been previously demonstrated (35), chronic DOX resulted in an increased number of lipid droplets in WT (1.75-fold, p<0.05), whereas there was no significant increase in lipid in β_2_-KO (Figure 3B). When we examined mice exposed to the lower dose sub-acute DOX (4 mg·kg^-1^ for 2 days), we found no difference in lipid droplets between WT and β_2_-KO (Supporting Information Figure S2).

**Figure 3.**
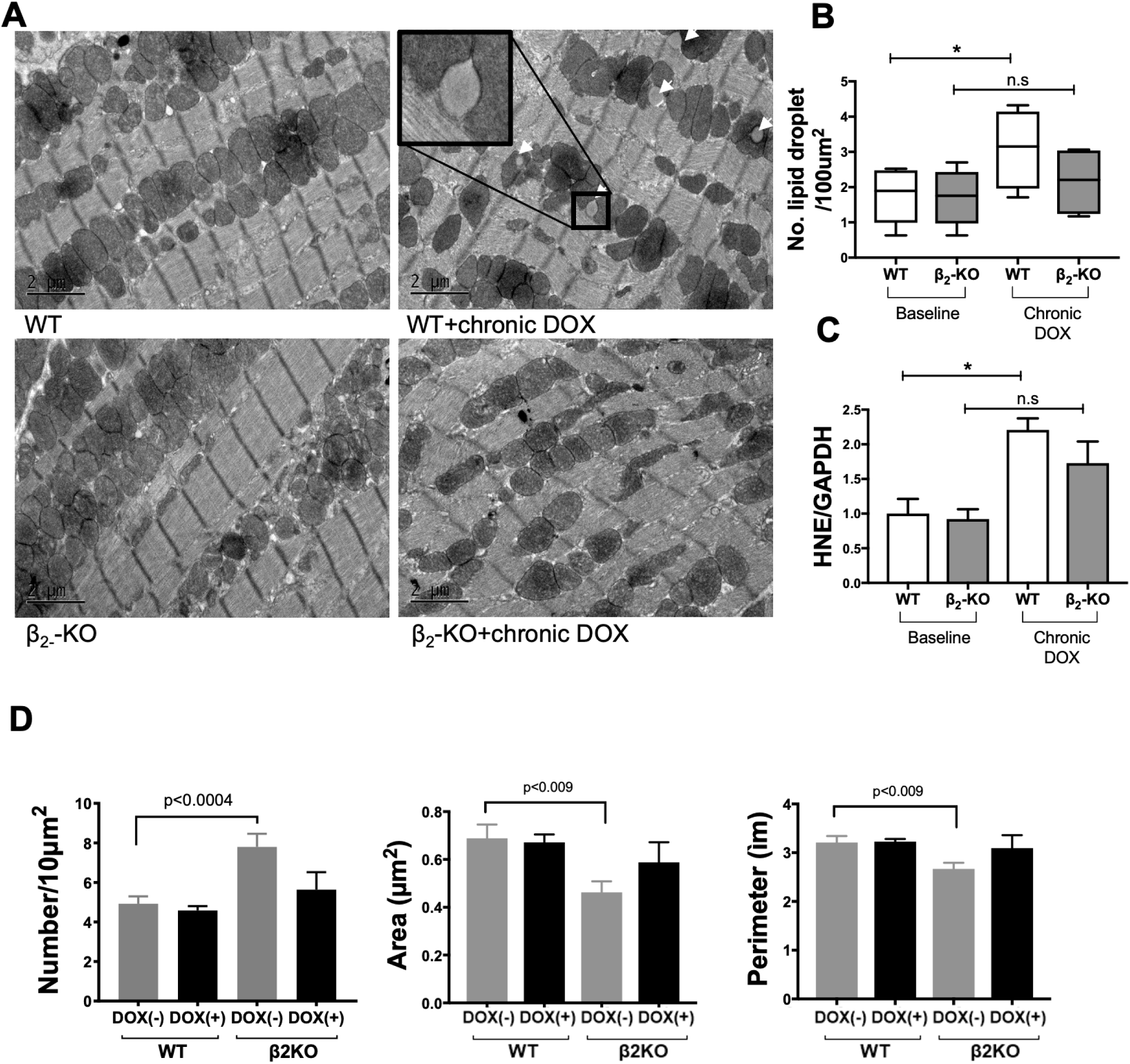
Deletion of the β_2_-AR reduces lipid accumulation and oxidative stress after chronic DOX. **A**. Electron micrographs from LV myocardium show that mitochondrial disarray occurs in both WT and β_2_-KO after chronic DOX, but increased lipid droplets (arrows) occur only in WT and not in the β_2_-KO. **B.** Numbers of lipid droplet were quantified in each condition. **C.** HNE staining (normalized by GAPDH) was significantly increased in WT with chronic DOX, but less increased in β_2_-KO, which failed to reach statistical significance. Data show mean ± SEM, * p<0.05 and n.s, not significant (one-way ANOVA with Tukey test). **D.** Differences in mitochondria morphology: number, area, and perimeter, were not evident after chronic DOX treatment between WT and β_2_-KO mice. β_2_-KO mice had evidence of mitochondrial fission at baseline (increased number of smaller mitochondria) as previously reported (34) but this difference resolved after chronic DOX.

Lipid droplets, as amphiphilic components, induce adjacent mitochondria membrane lipid peroxidation, which is one of the main mechanisms of doxorubicin toxicity in cardiomyocytes. Therefore, we assessed the level of lipid peroxidation using 4-hydroxynonenal (HNE) antibody staining. In WT after chronic DOX, there was a significant increase in HNE staining (2.2-fold, p<0.05). In contrast, the β_2_-KO had a lower level of lipid peroxidation than WT, which although trending higher, did not reach statistical significance compared to baseline (Figure 3C).

Differences in mitochondrial morphology (number, area, and perimeter) consistent with fission or fusion were not evident after chronic DOX treatment between WT and β_2_-KO mice (Figure 3D). However, mitochondrial size was smaller in the β_2_-KO at baseline.

### Differential gene expression during DOX treatment in WT and β_2_-KO mice

To better understand the cardioprotective mechanisms of the β_2_-KO during chronic DOX treatment, we undertook an unbiased transcriptome-wide analysis. Baseline (4 each/group) and chronic DOX (3 each/group) were analyzed. In addition, we used a subacute dose of DOX (4 mg·kg^-1^ for 2 days) to capture the early response to a lower dose of DOX (Figure 4A). As mentioned earlier, this was necessary given that the β2-knockouts all die within 15-30 minutes after acute DOX, too soon for reliable assessment of transcriptomic changes. We tested different sub-acute doses as follows: we have previously shown that a 15 mg·kg^-1^ dose results in 100% mortality in the β_2_-KO in 15-30 min. With a reduction in the acute DOX dose, mortality decreased (85% at 10 mg·kg^-1^; 63% at 8 mg·kg^-1^), but although the mortality rate went down, all β_2_-KO mice that died still did so within 15-30 min (8, 9). In contrast, a 4 mg·kg^-1^ dose resulted in mortality only after 4 days, therefore we used this subacute dose of DOX (4 mg·kg^-1^ for 2 days) to discern early (subacute) vs. late (chronic) gene changes in response to DOX. Differentially expressed genes (DEG) were identified from subacute and chronic groups after normalizing for baseline (subacute vs base, chronic vs base) or from chronic DOX after normalizing for subacute group (chronic vs subacute) in WT and β_2_-KO, respectively (Figure 4A). Each condition was also directly compared in β_2_-KO vs WT (β_2_-KO vs. WT panel, Figure 4A). Numbers of DEG from all comparisons are shown in bar graph format (cutoff is log2FC >0.4, adjusted p value <0.1).

**Figure 4.**
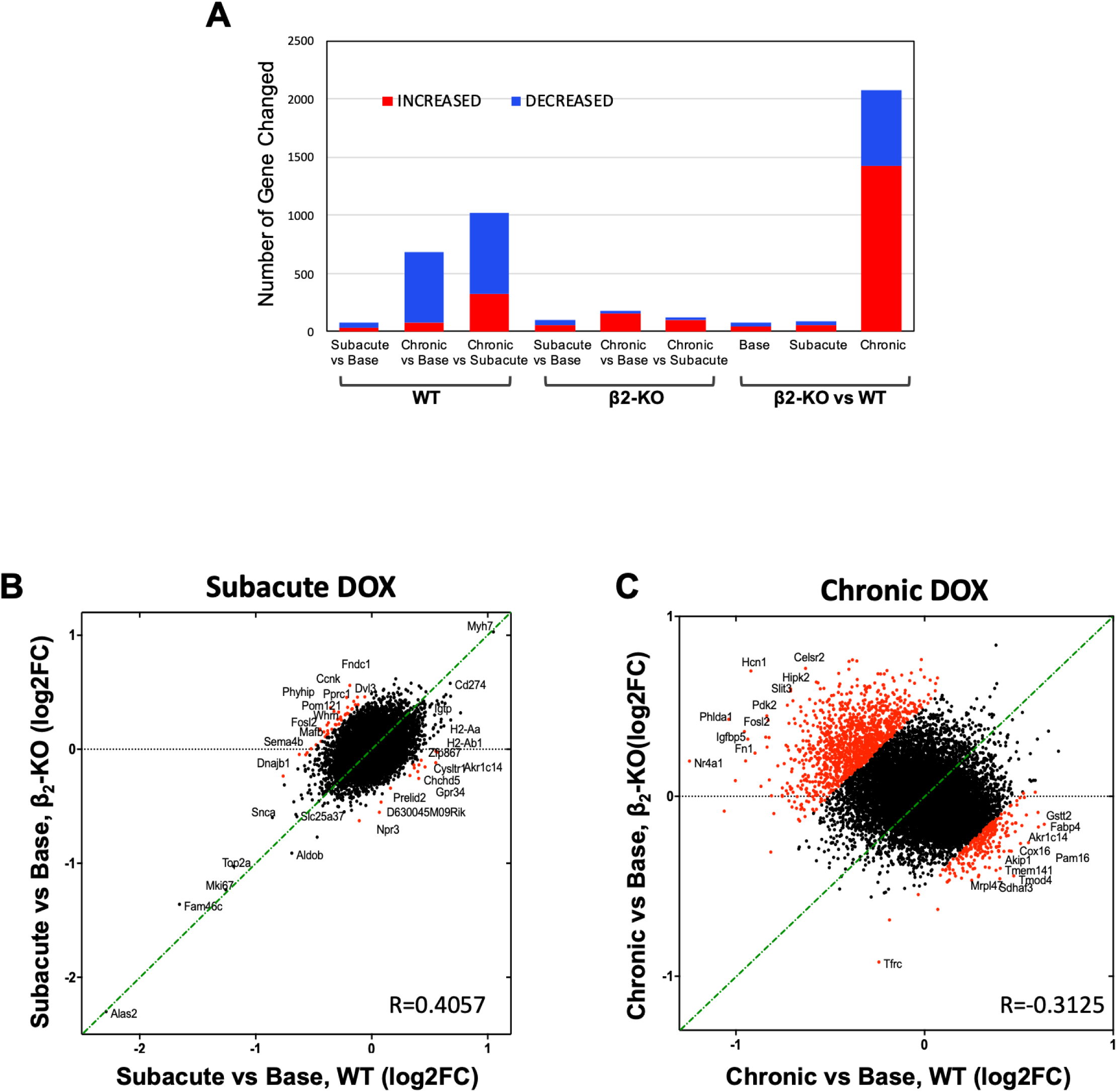
Differentially expressed gene (DEG) analysis in subacute and chronic DOX models. **A.** Stacked bar graph shows number of increased (red) and decreased (blue) DEGs under each condition. In the first two sections of this graph each group was compared to its baseline (subacute and chronic to baseline in WT and β_2_-KO; in the rightmost section, β_2_-KO was compared to WT, comparing each condition (baseline, subacute and chronic). Cut-off value is log2 fold change (log2FC) >0.4, adjusted p value (q value) <0.1. **B and C.** Scatter plots present comparison of the changes in WT and β_2_-KO mice during subacute (**B**) and chronic DOX (**C**). Fold changes were compared and the genes over 0.5 fold change difference are marked as red dots. Pearson correlation coefficients show relative relationship between groups. Subacute DOX has r=0.4057, and chronic DOX has r=-0.3125.

At baseline, deletion of the β_2_-AR altered transcription of only 74 genes compared to WT (42 upregulated and 32 downregulated, Supporting Information, Figure S3B. During subacute DOX, a total of 92 genes were differentially regulated, whereas during chronic DOX, a more dramatic 2077 genes were altered. 41 genes were differentially regulated in common between subacute and chronic DOX treatment in β2-KO vs WT (listed in Supporting Information, Figure S3A, B). Fibroblast growth factor 1(*FGF1*), ephrin type-B receptor 1(*EPHB1*), GTPases (rho GTPase-activating protein 6 (*ARHGAP6*) and ras-related protein Rab-27B (*RAB27B)* were all dramatically upregulated and phospholipid hydroperoxide glutathione peroxidase (*GPX4*) and NPC intracellular cholesterol transporter 1 (*NPC1*) were downregulated (Supporting Information, Figure S3C).

Chronic DOX treatment in WT mice changed significant numbers of genes compared to WT baseline, consistent with a previous report by Tokarka-Schlattner et al. (37), with most being downregulated (total 679 genes: 78 upregulated and 601 downregulated). In contrast, in β_2_-KO mice, a much smaller number of total genes were altered (total 182 genes) and the direction of their change was different: 158 upregulated and 24 downregulated. The most dramatic change was when we performed a direct comparison between β_2_-KO vs. WT after chronic DOX, increasing by 3-fold the number of altered genes (total 2075 genes: 1421 upregulated and 654 downregulated). These data suggested that the regulatory mechanisms induced by chronic DOX treatment are turned on/off in opposite directions in β_2_-KO vs. WT (Figure 4A).

Scatter plots present the pattern of responses during subacute (Figure 4B) and chronic DOX (Figure 4C) in both WT and β_2_-KO and highlight the distinctions between the two groups. Distribution of commonly changed genes are tightly aligned with the diagonal (green dashed line) and differentially regulated genes are divergent (red dots, >0.5 log2FC difference between β_2_-KO and WT). Subacute DOX treatment resulted in minimal changes in both groups of mice (74 in WT vs. baseline and 100 in β_2_-KO vs. baseline) which were similarly clustered in the center with no significant changes in either WT or β_2_-KO (Pearson’s correlation coefficient (R)=0.4057). Subacute DOX induces the common early responsive/compensatory pathway in both β_2_-KO and WT (20 genes in common) by controlling iron homeostasis in mitochondria [e.g. 5-aminolevulinate synthase, erythroid-specific, mitochondrial (*ALAS2*), rate-limiting heme biosynthesis, and mitoferrin-1 (*SCL25A37*) which works as an iron importer in mitochondria] and immune response [e.g. programmed cell death 1 ligand 1 (*CD274*), involved in T-cell proliferation and IFNG production, interferon gamma-induced GTPase (*IGTP*)] that are specific to subacute and not observed during chronic DOX (Supporting Information Figure S4A, C). Upstream regulator analysis using IPA predicts the commonly activated upstream regulators for immune response (interferon gamma (*IFNG*), signal transducer and activator of transcription 1 (*STAT1*), interleukin-21 (*IL21*)), and *TP53* shared in WT and β_2_-KO after subacute DOX, but also some unique or opposing factors in β_2_-KO (e.g. ERK1/2, BCL2/adenovirus E1B 19 kDa protein-interacting protein 3-like (*BNIP3L*), an apoptosis regulator) shown in Supporting Information Figure S4B.

In chronic DOX treatment, the scatter plot (Figure 4C) shows that a significant number of genes were deviated from the diagonal with negative correlation (R=-0.3125); red dots on the left side from the diagonal signify that many preserved or *downregulated* genes in WT were either preserved or *increased* in the β_2_-KO. Right side red dots signify the opposite effect.

### During chronic DOX, β_2_-KO mice show preservation of cardiac contractile genes and enhanced energy metabolism-related genes: protective mechanisms during chronic oxidative stress

To provide further insight into the protective mechanism operative in β_2_-KO during chronic DOX treatment, we examined DEGs in the chronic DOX group. Although we found 41 genes with altered regulation common to both β_2_-KO and WT after chronic DOX (Figure 5A), it is notable that for 39 out of these 41 genes the *directionality* of altered gene regulation was totally opposite between the two groups: upregulated in β_2_-KO and downregulated in WT. These included myocyte-specific enhancer factor 2D (*MEF2D),* cyclic AMP-dependent transcription factor *(ATF7),* acetyl-CoA carboxylase 2 *(ACACB),* insulin receptor substrate 1*(IRS1)*, and pro-low density lipoprotein receptor-related protein 1 (*LRP1)* (listed in Supporting Information Figure S5A).

**Figure 5.**
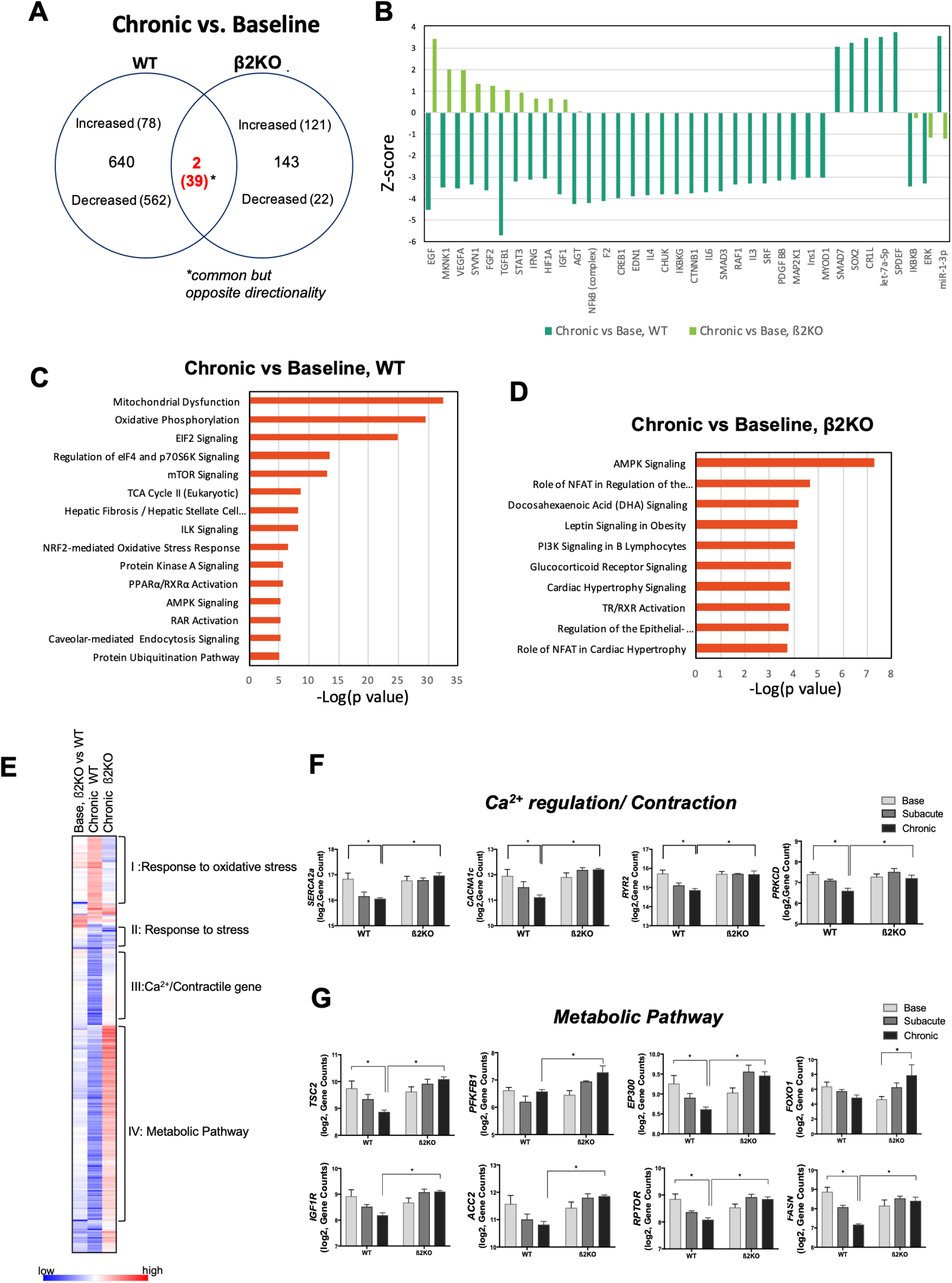
Modulation of gene expression in WT and β_2_-KO during chronic DOX treatment: The β_2_-KO was protected from chronic DOX-induced dysregulation in Ca^2+^/contractile and metabolic pathways. **A.** Venn diagram presents the number of altered genes in WT and β_2_-KO. Critically, among 41 common altered genes, 39 are downregulated in WT but upregulated in β_2_-KO, * show DEGs in common but directionality opposite, only 2 genes were changed in the same direction. **B.** Upstream regulators analyzed by IPA showed that WT and β_2_-KO shared high ranked upstream regulators but in opposite direction (the correlation in z-score is opposite): EGF, VEGFA, FGF2, and TGFB1. **C** and **D**. Significantly altered canonical pathways in chronic DOX vs. baseline in WT and β_2_-KO. **E**. The “organism survival” related gene set showed significant differences in directionality of altered expression between WT and the β_2_-KO. This gene set was extracted from IPA bio-functional analysis and compared at baseline between WT and β_2_-KO (Base, β_2_-KO vs WT), between baseline and chronic DOX in the WT (Chronic WT) and between baseline and chronic DOX in the β_2_-KO (Chronic β_2_-KO). Whereas there were minimal changes between WT and β_2_-KO at baseline, in response to chronic DOX, WT showed decreased expression in categories of Ca^2+^ regulation/contractile genes and metabolic/energetic pathway genes. In contrast, the β_2_-KO showed no changes in Ca^2+^ regulation/contractile genes, and increased expression in metabolic/energetic pathway genes. Similar changes in stress genes were seen between WT and β2-KO in response to DOX. The second set of comparisons is between the WT and β_2_-KO after chronic DOX**. F**. Contractile machinery genes (Ca^2+^ handling/sarcomeric molecules) were downregulated in WT and preserved in β2-KO mice after chronic DOX. **G**. Energy metabolism related genes were downregulated in WT and upregulated in β2-KO mice after chronic DOX. Data show mean ± SEM of Log2, gene counts (HTSeq-count), * *p*<0.05 and n.s, not significant (one-way ANOVA with Tukey test).

We performed analysis for biological pathways and upstream regulator pathways using IPA (Figure 5C, D). As shown previously, our study confirmed that a main target of DOX toxicity in cardiomyocytes is the mitochondria (38). In WT after DOX, mitochondrial dysfunction and oxidative phosphorylation were the most significantly affected canonical pathways. In addition, oxidative stress response and metabolic pathways (mTOR/S6K signaling/AMPK signaling/TCA cycle) were significantly altered in WT (Figure 5C). In contrast, in β_2_-KO after chronic DOX, the most affected canonical pathways are AMPK signaling/PI3K and NFAT regulating signaling/cardiac hypertrophy (Figure 5D), independent of directionality. We then analyzed pathways with emphasis on directionality. Upstream regulator analysis in chronic DOX groups also revealed that the most common and highest ranked regulators were negatively correlated in WT but positively correlated in β_2_-KO (e.g. epidermal growth factor (EGF), MAP kinase-interacting serine/threonine-protein kinase 1 (MKNK1), vascular endothelial growth factor A (VEGFA), transforming growth factor beta-1 (TGFB1) (Figure 5B).

The most differentially regulated bio-function pathway between WT and β_2_-KO was “mortality in organism” (z score=12.55, p value =5.62E-31 in WT, z score=-7.5, p value=1.15E-13 in β_2_-KO). This pathway of 355 genes was extracted and compared (Figure 5E, F, G) revealing some of the more marked changes between these two groups. During baseline conditions, β_2_-KO mice express similar levels of these genes to WT mice (baseline, β_2_-KO vs. WT), however, after chronic DOX, dramatic differences develop. Based on the pattern of changes in WT and β_2_-KO, 4 groups are categorized:

(I) response to oxidative stress; (II) Response to stress; (III) Ca^2+^/contractile genes; and (IV) Metabolic pathways. In WT mice after chronic DOX, Ca^2+^ handling/sarcomeric contraction genes (Group III), many of which have been shown to be altered in heart failure, were downregulated (Figure 5F) including L-type calcium channel (*CACNA1C*), ryanodine receptor 2 (*RYR2*), sarcoplasmic/endoplasmic reticulum calcium ATPase 2 (*ATP2A2*), protein kinase delta c (*PRKCD*), sodium channel (*SCN5A*), myosin binding protein 3 (*MYBPC3*), titin (*TTN*), myosin heavy chain 6 and 7 (MYH6/7), alpha cardiac muscle actin 1 *(ACTC1*). Transcription factors that regulate cardiac gene expression were also downregulated (*GATA4,* Heart and neural crest derivatives-expressed protein 2 *(HAND2),* myocardin *(MYOCD)* and *MEF2D*). In contrast, in the β_2_-KO, gene expression in Ca^2+^ handling/sarcomeric contraction (group III) was preserved (Figure 5E), consistent with the delayed cardiac dysfunction in the β_2_-KO. Genes in group I (Response to Oxidative Stress), which were mostly upregulated in WT, were unchanged or slightly decreased in the β_2_-KO, mostly in mitochondrial and oxidative stress response pathways (perosiredoxin1 (*PRDX1*), thioredoxin (*TXN),* superoxide dismutase 1 (*SOD1)*, glutathione peroxidase 4 (*GPX4)*, NADH quinone dehydrogenase 1 (*NQO1)*, isocitrate dehydrogenase 1, cytosolic *(IDH1)*). Although most genes in the stress responsive system were differentially regulated in WT vs. the β_2_-KO, some were commonly downregulated (heat shock protein beta-8 *(HSPB8),* peptidyl-prolyl cis-trans isomerase *(FKBP4),* dnaJ homolog subfamily B member 1*(DNAJB1),* aquaporin-1*(AQP1)*) in both β_2_-KO and WT with chronic DOX (group II, Figure 5E).

Metabolism-related genes (group IV, Figure 5E) also showed dramatic differences between WT (downregulated) and the β_2_-KO (upregulated) with chronic DOX. The AMPK signaling pathway was the most altered in this group (Supporting Information Figure S5B). In WT we observed downregulation of a set of AMPK-associated genes (serine/threonine-protein kinase ULK1(*ULK1*), CREB-binding protein (*CREBBP*), histone acetyltransferase p300 (*EP300*), fibroblast growth factor receptor 1(*FGFR1*), insulin receptor substrate 2(*IRS2*), forkhead box protein O1(*FOXO1*), tuberin (*TSC2*), fatty acid synthase (*FASN*), insulin-like growth factor 1 receptor (*IGF1R*)) and AMPK-regulated transcription factors, including PPARG coactivator 1 alpha (*PPARGC1a*). In contrast, these were all upregulated in the β_2_-KO, suggesting that chronic metabolic adjustments related to ablation of β_2_-AR signaling may represent one of the compensatory survival responses during chronic oxidative stress such as chronic DOX (Figure 5G).

### Cardioprotective/compensatory pathways that are activated early during DOX treatment are sustained in the β_2_-KO, but not in WT

The distinct transcriptional changes in WT and β_2_-KO during chronic DOX are compared with the pattern of gene changes during sub-acute DOX treatment. Interestingly, Figure 4A shows that in WT, the number of DEGs between chronic vs. subacute DOX (total 1019 genes: 328 upregulated and 691 downregulated) is larger than that between chronic DOX vs. baseline (total 691), suggesting that early compensatory responses during subacute DOX were transitory in the WT, leading to differential gene changes that may have contributed to their earlier decompensation with chronic exposure to DOX. In contrast, in the β_2_-KO, the number of DEGs between chronic vs. subacute DOX (total 123 genes: 106 upregulated and 12 downregulated), is smaller than that in WT and also somewhat smaller than between chronic DOX vs. baseline (182), suggesting that the compensatory response that was turned on early after DOX was sustained. Scatter plots of WT and β_2_-KO (Figure 6A and B) also show a greater correlation between subacute and chronic DOX in the β_2_-KO compared to WT (R=0.4683 in β_2_-KO vs R=0.2707 in WT).

**Figure 6.**
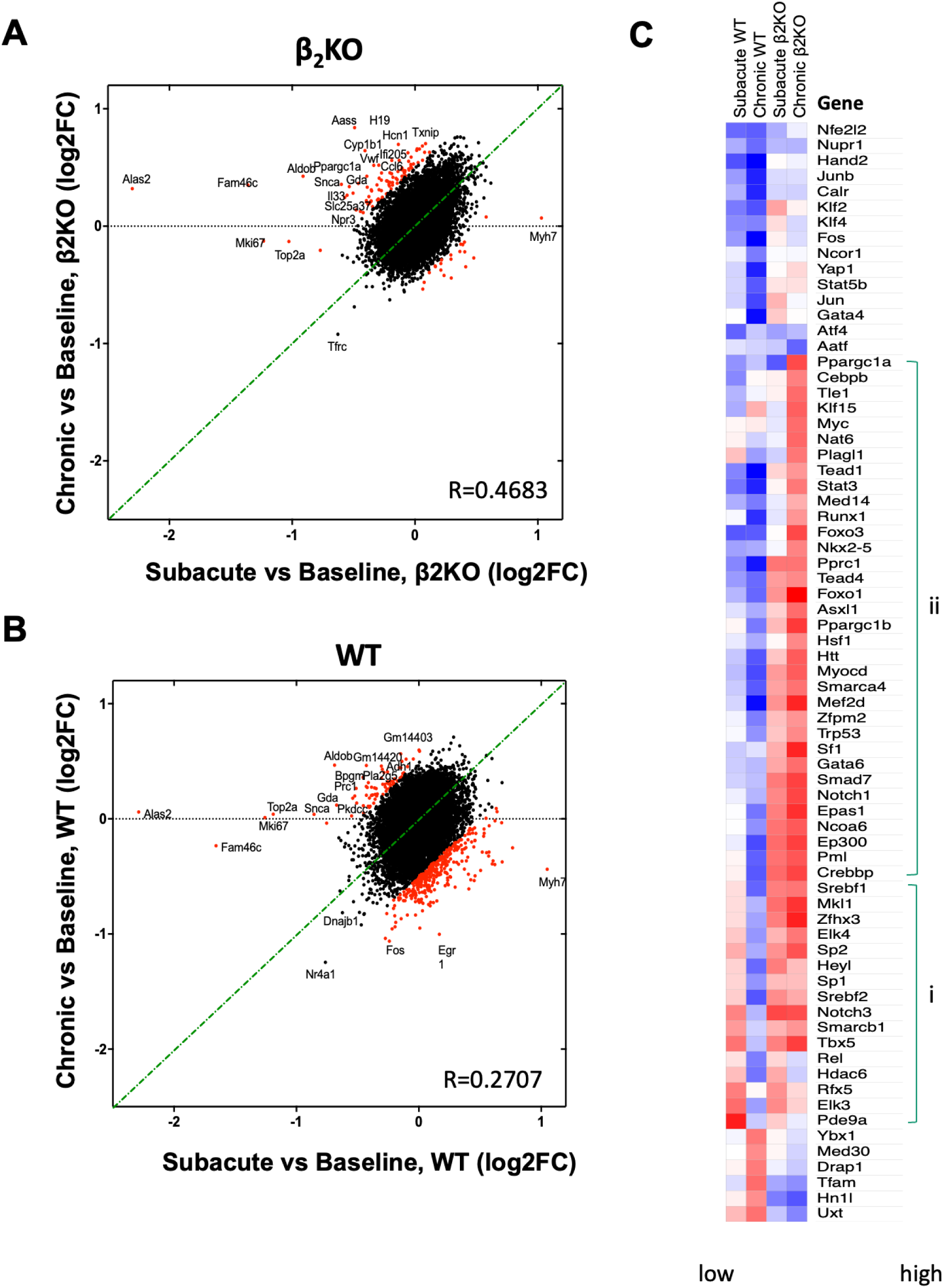
Comparison between subacute (early) and chronic (late) response in WT vs. β_2_-KO. Scatter plots in **(A)** β_2_-KO and **(B)** WT compare subacute vs. chronic responses. There is better correlation between subacute and chronic DOX gene expression in the β_2_-KO, and more differentially downregulated genes in the WT. **(C)** Comparison of transcription factor expression (subacute vs. chronic DOX in WT and in β_2_-KO). In WT, TFs are mostly downregulated with chronic DOX, whereas in the β_2_-KO many of these key regulatory TFs are increased.

To develop further insight into differential on/off regulation of the cardioprotective mechanism during the early and late response to DOX in the β_2_-KO, we compared the expression pattern of transcription factors (TFs) that are part of the earliest response to stress and propagate their effect through downstream gene regulation. We extracted the most significantly altered TFs (71) and compared them between each condition (subacute and chronic DOX in WT or β_2_-KO). Changes of TF expression are shown by color index to enhance visualization of the trend (Figure 6C). Interestingly, from the panel of compared TFs, similarity clustering shows that with chronic DOX in WT, TFs are mostly downregulated, whereas in the β_2_-KO, the key regulatory TFs are activated and most of the increased TFs are also upregulated during subacute DOX, including cardiac TFs (*MYOCD, MEF2D*), metabolism related TFs (*FOXO1, PPARGC1a/b, EP300, CREBBP, NCOA6, EPAS1* Sterol regulatory element-binding protein 1/2 (*SREBF1/2*)), and SWI/SNF chromatin remodeling complex (*SMARCA4, SMARCB1*)). These data support the paradigm that TFs may be early markers for detecting either detrimental or beneficial response to chronic cardiotoxic stress.

### CaMKII and AMPK signaling are differentially regulated in β_2_-KO vs WT both at baseline and after DOX

In the absence of β_2_-AR signaling, we have previously shown that basal cytosolic Ca^2+^ is increased (6), due to removal of β_2_-AR-G_i_ inhibition of β_1_-AR-G_s_ signaling. We have also shown that this increase in Ca^2+^ plays a cardioprotective role under certain stress conditions such as pressure-overload induced cardiomyopathy and MLP-KO mediated dilated cardiomyopathy (14). These data, in combination with our current finding of differential expression of Ca^2+^ regulatory genes in the β2-KO with chronic DOX, led us to question whether ablation of the β_2_-AR could mediate a beneficial role in chronic DOX via alterations in Ca^2+^, similar to the above models. The Ca^2+^-regulated kinase CaMKII is one potential effector of β-AR signaling, having been shown to play a major role in cardiac remodeling during multiple stress conditions (39). We found that, at baseline, CaMKII phosphorylation is increased in the β_2_-KO without an alteration in total CaMKII expression. This increase persisted with chronic DOX (Figure 7). Among the 3 major cardiac isoforms of CaMKIIδ (a, b and c), CaMKIIδb phosphorylation was most significantly increased in the β_2_-KO compared with the other isoforms (a and c) (Supporting Information Figure S6).

**Figure 7.**
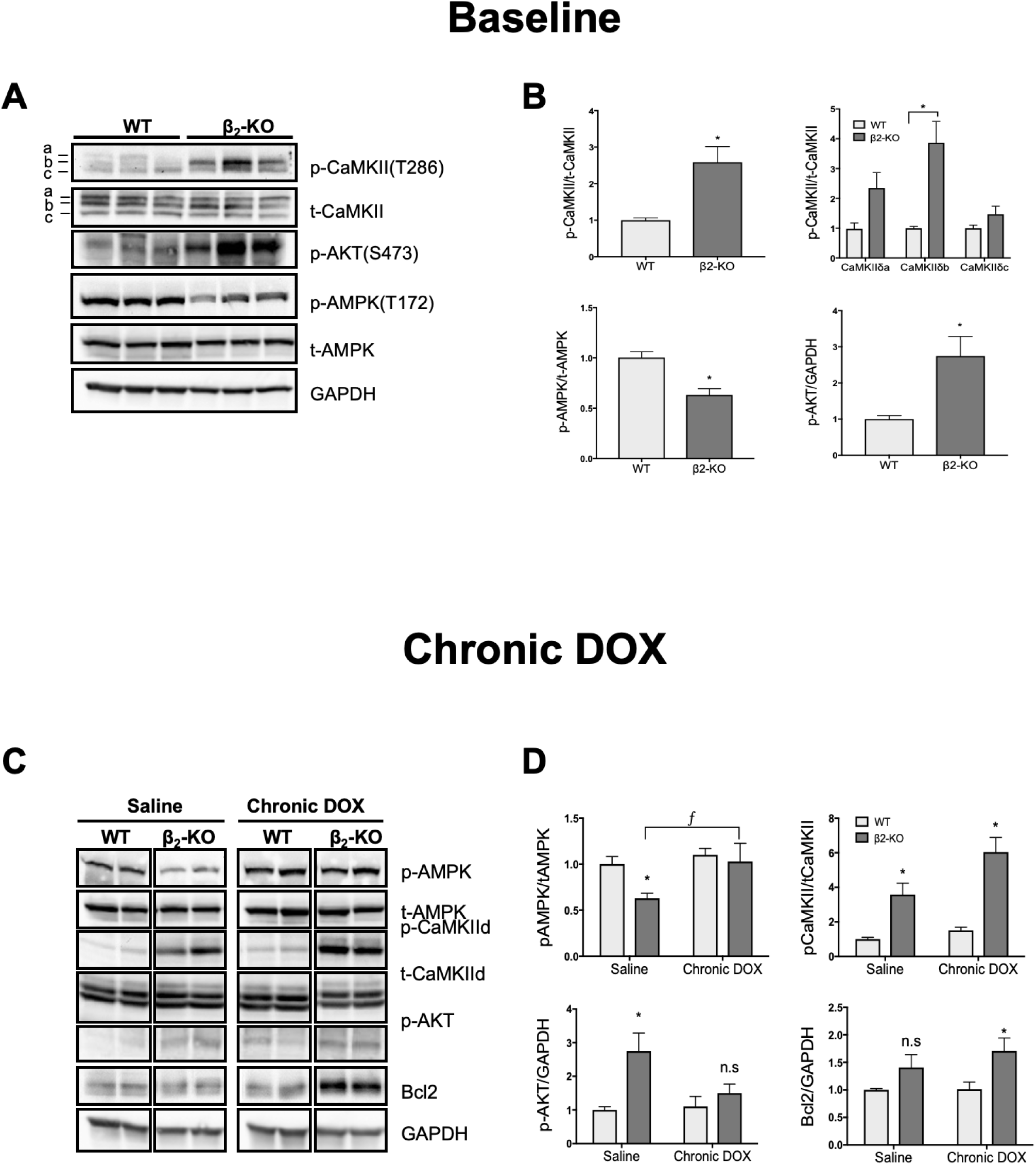
CaMKII, AMPK and AKT signaling are differently regulated in β_2_-KO vs. WT. **A**. Western blots were performed to detect expression and phosphorylation profiles of kinases (Ca^2+^/calmodulin kinase II (CaMKII), AMPK, and AKT)) at baseline; CaMKIIδ isoforms (a, b and c) were separated by molecular weight. **B**. Blots were quantified and normalized by loading control (either total protein for each phosphorylated form or GAPDH) **C**. Western blots were performed to detect expression and phosphorylation profiles of kinases (Ca^2+^/calmodulin kinase II (CaMKII), AMPK, and AKT) and cardioprotective molecules such as Bcl2 after chronic DOX. **D**. Blots were quantified and normalized by loading control (either total protein for each phosphorylated form or GAPDH). Most notably, phosphorylation of CaMKII was increased in the β2-KO vs WT at baseline, and persisted with chronic DOX, but did not increase further. This increase was most notable for CaMKIIδb (see Figure S6). Data show mean ± SEM, * p<0.05 and n.s, not significant, β2-KO vs. WT (one-way ANOVA with Tukey test).

Our transcriptome analysis showed that metabolic pathways are most affected in the β_2_-AR during chronic DOX, including the important AMPK signaling pathway. The level of activation of AMPK was tested by assaying AMPK phosphorylation (Figure 7). The β_2_-KO at baseline shows a marked 40% downregulation of phospho-AMPK vs. WT. In contrast, chronic DOX enhances AMPK phosphorylation in the β_2_-KO but not in WT, suggesting that the increased capacity to activate AMPK signaling may play a protective role in the β_2_-KO during chronic stress induced by DOX, even though the levels of AMPK phosphorylation after chronic DOX in WT and β_2_-KO were similar.

## DISCUSSION

Signaling through β1 and β2-ARs is a principal regulator of normal cardiac inotropy and chronotropy but also mediates adverse remodeling in heart failure associated with sustained sympathetic activation (1). β_1_-ARs are subtype-selectively downregulated in heart failure and thus drugs targeting β-ARs (either β_1_-specific blockers such as metoprolol or β_1_β_2_ blockers such as carvedilol) are standard of care and highly effective therapies to reduce sympathetic-induced injury and restore normal β-AR signaling homeostasis (2, 3). Prior research suggests that, in addition to its effects on inotropy and chronotropy, the β_1_-AR predominantly induces cardiotoxic signaling, whereas the β_2_-AR induces direct cardioprotective signaling, as well as opposing β_1_-mediated cardiotoxic signaling through activation of Gi and by crosstalk with regulators such as CaMKII, MAPKs and PI3K-Akt (4–7). However, there is conflicting data on the role of β_2_-ARs, differing dependent on the specific stressor and the duration of exposure to that stressor. The goal of this study was to understand the differential cardioprotective/cardiotoxic effects of β_2_-AR signaling dependent on timing (acute vs. chronic) during oxidative cardiac stress.

We chose a model of oxidative stress, the anthracycline anti-cancer drug doxorubicin (DOX). One of the oldest chemotherapeutic agents, DOX is associated with clinical cardiotoxicity, and has long been utilized as an animal model of oxidative-stress cardiotoxicity. DOX cardiotoxicity can be acute, during or shortly after its administration, or chronic, where cardiotoxicity occurs later, after the last dose of administration, and in patients, even as long as years afterwards. Clinically, this toxicity is the major side effect that limits its clinical use (40). The mechanisms underlying DOX-induced cardiotoxicity include reactive oxygen species (ROS) generation, lipid peroxidation and energetic imbalance caused by mitochondrial dysfunction (41, 42), alteration in iron homeostasis, calcium dysregulation, injury to cardiac myofilaments as well as dsDNA breakage through topoisomerase IIβ (43, 44).

There have been multiple previous studies investigating the role of pathways downstream of β-ARs, specifically the MAPKs, in mediating DOX toxicity. *In vitro*, the anthracycline daunomycin activates ERK1/2, p38 MAPK and JNK in neonatal rat cardiomyocytes through reactive oxygen species and Ca^2+^, with data suggesting that ERK1/2 protects cardiomyocytes from apoptosis, whereas p38 MAPK is involved in the induction of apoptosis (45). DOX administration to adult rat cardiomyocytes increases ROS levels and the activation of stress-induced proteins p53, p38 and JNK MAPKs, inducing autophagy and apoptosis (46). We have previously shown that JNK is differentially activated in β_2_-KO neonatal cardiomyocytes after DOX. JNK inhibition increases toxicity, suggesting that JNK is acting in a cardioprotective role in our model (9). However in H9c2 cardiac muscle cells, JNK activity has been associated with DOX-induced apoptosis (47). Differential phosphorylation of downstream signaling associated with JNK could be involved in its intricate regulation of the balance between cardiotoxicity and cardioprotection (9). *In vivo* studies have also focused on the role of MAPKs in DOX cardiotoxicity. Phosphorylation of ERK1/2, in the early stage of DOX induced cardiomyopathy in rats, shows a biphasic response, with a massive 5-fold increase at 4 h, followed by a decrease to 67% 3 weeks after the last injection. Phosphorylation of p38 and JNK steadily increases in the first 24 h and after 3 weeks (48).

We have shown that a differential activation of MAPK isoforms, previously shown *in vitro* to regulate β-agonist as well as DOX cardiotoxicity, appears to play a role in mediating the differential temporal effects of these β-adrenergic receptor subtypes in vivo. Although MAPKs are highly activated following acute DOX treatment in β2-KO mice compared to WT, with p38 the most differentially upregulated (8), we did not observe activation of any MAPK family member (p38, ERK1/2, JNK) at 7 weeks following chronic low-dose DOX treatment, at which time β2-KOs showed a survival benefit. The absence of the β2-AR in the context of high levels of stress induced by an acute high dose of DOX activates the MAPK response, with p38 increased by 20 fold and 100% mortality (8); however, in the absence of the β2-AR, this stress response is lost under chronic DOX stimulation. It is possible that other signaling pathways may be involved in the inactivation of MAPK signaling with increased duration of DOX administration, however this decrease may partially be responsible for the cardioprotection conferred by the absence of the β2-AR in the context of chronic DOX stimulation.

We have previously shown that the MAPK inhibitor SB-203580 rescues β2-KO mice from acute toxicity (8); furthermore, studies with transgenic mice with cardiac-specific expression of a dominant-negative mutant form of p38α suggest that p38α MAPK may play a role in the regulation of cardiac function, oxidative stress, and inflammatory and apoptotic mediators in the heart after DOX administration (49). In addition, deletion of p38δ and p38γ/δ has been shown to be cardioprotective (50). This opposite effect of β2-AR signaling on the activation of MAPK suggests that its signaling may function in a cell type-dependent and stimulus-dependent manner in regulating the balance between cardiotoxicity and cardioprotection.

Chronic treatment with DOX has been shown to induce several morphological alterations, including disarray and structural alterations in mitochondria and accumulation of lipids (35, 36). Cardiac mitochondria are a primary target of DOX toxicity and mediate the generation of highly reactive free radical species of oxygen (ROS), accumulation of iron and alteration in metabolic processes (51). We have previously shown that β_2_-KO cardiomyocytes show mitochondrial fission: smaller size, perimeter and surface area and larger number of mitochondria compared to WT (34). Similarly, in acute DOX (15 mg·kg^-1^ for 24 h) given a p38 inhibitor, β_2_-KO mice develop exaggerated mitochondrial disruption compared to WT (6). In the chronic DOX setting, the β_2_-KO shows the same degree of structural disarray of mitochondria as the WT but with significantly attenuated lipid droplet formation and lipid peroxidation.

We found that CaMKII phosphorylation is increased in the β_2_-KO without an alteration in total CaMKII expression and this increase persisted with chronic DOX. At first glance, this would suggest increased activation of adverse remodeling signaling. In adult rat cardiomyocytes DOX exposure induces CaMKII-dependent SR Ca^2+^ leakage, contributing to impaired cellular [Ca^2+^]_(i)_ homeostasis (52). In neonatal rat cardiomyocytes DOX has been shown to activate CaMKII promoting apoptosis (53). However, different cardiac isoforms of CaMKIIδ have been shown to regulate opposing cardiotoxic/cardioprotective effects. In the heart, CaMKIIδB and CaMKIIδC, localize in nuclear and cytosolic compartments, respectively. Activation of CaMKIIδC plays a role in cardiomyocyte apoptosis and has been associated with increased mitochondrial cytochrome c release (54). In contrast, overexpression of CaMKIIδB protected cardiomyocytes against oxidative stress, hypoxia, and angiotensin II induced apoptosis (55). Among the 3 major cardiac isoforms of CaMKIIδ (a, b and c), CaMKIIδb phosphorylation, the isoform associated with *cardioprotection*, is most significantly increased in the β_2_-KO compared with the other isoforms. Although this baseline increase of CaMKIIδb phosphorylation would help to protect against the acute cardiotoxicity observed in the β_2_-KO mice we speculate that it’s not enough to counteract the severe overload in acute DOX toxicity; however, in chronic DOX this increased phosphorylation of the cardioprotective isoform could be one mechanism to ameliorate toxicity.

To better elucidate the molecular basis of β_2_-AR signaling switching from being cardioprotective to cardiotoxic, we performed a serial transcriptomic analysis using RNA-seq on hearts from our mouse model. These data show that DOX treatment in WT mice results in a much larger number of gene expression changes compared to β_2_-KO mice. The most dramatic change was seen when comparing β_2_-KO vs WT after chronic DOX, suggesting the regulatory mechanisms induced by chronic DOX treatment are turned on/off in opposite directions in β_2_-KO vs. WT.

Further analysis of our data using IPA bio-functional analysis showed significantly altered canonical pathways in WT related to mitochondrial dysfunction and oxidative phosphorylation; whereas in β_2_-KO altered pathways include AMPK signaling/PI3K and NFAT regulating signaling/cardiac hypertrophy. By categorizing these sets of differentially expressed genes we found that the “organism survival” related genes and “contractile machinery” genes were most downregulated in WT after DOX. One example, RYR2 channels, are associated with many cellular functions, including mitochondrial metabolism, gene expression and cell survival, in addition to their role in calcium flux and cardiomyocyte contraction (56). Calcium flow through L-type calcium channels (*CACNA1C*) is an essential step in initiating the signaling cascade that leads to cardiac muscle contraction (57). Another study reported that SERCA2 knockout heterozygous hearts showed impaired cardiac function *in viv*o (58). Critically, all of those genes were preserved in the β_2_-KO, consistent with the enhanced survival and delayed cardiac dysfunction in β_2_-KO mice. We also compared the expression pattern of transcription factors (TFs) that are part of the earliest response to stress to develop further insight into differential on/off regulation of the cardioprotective mechanisms during the early and late response to DOX in the β_2_-KO. Several key regulatory TFs that are most increased in chronic DOX are also upregulated during subacute DOX in the β_2_-KO. These results provide evidence that TFs may be early markers for detecting either detrimental or beneficial responses to chronic cardiotoxic stress.

Impairment of cardiac bioenergetics is critically important in the pathogenesis of DOX-induced cardiotoxicity and has emerged as a potential therapeutic target (59). DOX reduces basal high energy phosphate levels and oxidative capacity of mitochondria, as well as induces defects in the AMPK signaling pathway (60, 61). AMPK is known to respond to minor changes in the AMP/ATP ratio as well as other physiological and pathological stimulation (such as exercise, oxidative stress, hypoxia, and glucose deprivation) (62). It is thus worthwhile to also focus on the energy metabolism-related genes which are dramatically *downregulated* in the WT with chronic DOX, whereas these were *upregulated* in the β_2_-KO with chronic DOX. AMPK signaling was the most altered of the metabolic pathways. AMPK, highly conserved across animal species, is an energy sensor and a cellular master regulator of energy homeostasis, autophagy and fibrosis (63–65). AMPK is activated upon phosphorylation of its α1, α2 and β1 subunits, and subsequently phosphorylates a multitude of downstream effectors (66). Specific AMPK agonists decrease cardiac fibrosis and increase cardiac ejection fraction, inhibit fatty acid synthesis and protect against ischemia-reperfusion injury in mice (67). In organs with high energy demands such as the heart, AMPK signaling activates both glucose uptake and glycolysis and promotes catabolic pathways as well as an increase in fatty acid oxidation (68–70). AMPK activation plays a beneficial role in cardiovascular protection and has been proposed as a therapeutic target in heart failure (71, 72).

AMPK sits at the control point of many mechanisms shown to be involved in doxorubicin cardiotoxicity. Preclinical studies showed that AMPK-activating agents may play a cardioprotective role in models of DOX cardiotoxicity (73). We found that altered phosphorylation of AMPK at baseline and during chronic stress may play an important cardioprotective role in the β_2_-KO. The absence of β_2_-AR signaling at baseline significantly decreased phosphorylated-AMPK vs. WT. There have been several studies suggesting an interaction between β-ARs and AMPK signaling. In a high-epinephrine rat model, the β-AR-AMPK pathway has been shown to phosphorylate acetyl-CoA carboxylase, reducing abdominal visceral fat accumulation and increasing insulin sensitivity (74). In addition, in studies where β-AR over-activation induces fibrosis, a decrease in AMPK activity, which further increases β-arrestin 1, has been postulated as a central mechanism of β-AR cell injury. The AMPK activator metformin has also been shown to decrease β-arrestin 1 expression and attenuate cardiac fibrosis (75). Carvedilol, a third generation β-blocker, protects the heart from ischemia-reperfusion injury in part through activation of the AMPK-acetyl-CoA carboxylase signaling pathway (76). Interestingly, β_2_-ARs have been shown to directly regulate metabolic signaling and function to induce the phosphorylation of AMPKα^T172^ activate PKB, and thereby modulate energy metabolism in rat ventricular muscle (77). In our current study, chronic DOX triggers a variety of cellular cascade reactions, especially enhanced AMPK phosphorylation in the β_2_-KO, but critically, not in the WT. Ferroptosis, a form of regulated cell death induced by iron-dependent lipid peroxidation can be inhibited by treatments that induce or mimic energy stress. Inactivation of AMPK largely abolishes the protective effects of energy stress on ferroptosis (78). Thus, the increased capacity to activate AMPK signaling only in the β_2_-AR exerts potentially protective effects during chronic oxidative stress such as occurs with chronic DOX.

AMPK is known to respond to minor changes in the AMP/ATP ratio. The ability of AMPK to reprogram metabolism is highly relevant for the understanding of chronic metabolic adjustments related to ablation of β_2_-AR signaling and their compensatory survival response. The examination of novel mechanisms of AMPK regulation and the identification of sets of AMPK-associated genes helped us to understand how β_2_-AR signaling switches from being cardioprotective in acute stress to cardiotoxic during chronic stress. For example, EP300 is an AMPK-associated gene which plays a role in regulating cell growth and maturation and is critical for normal development. Fibroblast growth factor receptor 1 (FGFR1) elicits cellular responses by activating signaling pathways, i.e. phospholipase C/PI3K/AKT; ras subfamily/ERK; protein kinase C, IP3-induced raising of cytosolic Ca^2+^, and Ca^2+^/calmodulin-activated elements and pathways (79). PPARG coactivator 1 alpha (PPARGC1α) encodes PGC-1α protein which provides a direct link between external physiological stimuli and the regulation of mitochondrial biogenesis (80). Forehead box protein O1(FOXO1) plays an important role in cellular proliferation, metabolism, inflammation, differentiation, and stress resistance (81). All of these genes are categorized in group IV: metabolic pathway (Figure 5D). Critically, these AMPK-associated genes were all *downregulated* in WT but *upregulated* in the β_2_-KO, suggesting that chronic metabolic adjustments related to ablation of β_2_-AR signaling may mediate a compensatory survival response following chronic DOX treatment.

Our study has several limitations: first, although we wanted to compare the effect of β*2*-AR regulation of signaling in acute DOX and chronic DOX we were able to do so only in the more canonical signaling pathways and their associated posttranslational modifications which can occur quite rapidly; we were not able to compare changes in gene expression directly due to the severity of acute DOX cardiotoxicity in the β*2*-KO, resulting in 100% mortality within 15-30 min, a time too short for meaningful analysis of changes in gene expression. We adopted a compromise approach to analyze these changes using a subacute dose of DOX to capture the early response in the β2-KO to DOX. Our findings showed dramatic acute vs. chronic differences in gene expression, but need to be placed in context of this limitation. Second, the comparison between acute vs chronic DOX is somewhat limited by the choice of doses we used; we could have studied additional doses to more completely explore dose-effect, rather than time-effect. Despite this limitation, we uncovered key alterations in gene expression and cell signaling that provide putative mechanisms for this unique time-dependent switch of signaling regulated by the β2-receptor in cardiotoxicity. We always controlled these experiments by comparing the β2-KO to WT under the same conditions. Future studies can supplement these findings using multiple dosing regimens, however, our fundamental finding regarding the dramatic difference in acute vs. chronic response to DOX in the β2-KO remains intact. Third, while we observed an extended survival in both β1-KO and β2-KO mice after chronic DOX we focused on the changes of β_2_-ARs, since it was only in the β2-KO where there was a time-dependent differential response. We recognize that since multiple pathways are involved in this β_2_-AR homeostatic switch, there is also potential for multiple signaling crosstalk between these pathways. Finally, we have not definitively proven the role of any specific pathway in this temporal toxicity switch by performing knockout or overexpression studies, but have provided confirming evidence by citing previous studies where available. We believe that trying to provide this level of mechanistic confirmation would be beyond the scope of this initial discovery study, but worth pursuing in follow-up studies.

In conclusion, β_2_-ARs act as a key homeostatic switch playing critical and differential cardiotoxic/cardioprotective roles in the context of various stress conditions. When combined with *<u>high level, acute oxidative stresses</u>* such as acute DOX or myocardial infarction (11), β_2_-AR signaling is cardioprotective, so that blocking or deleting the β*2*-AR results in deleterious effects. In line with this paradigm, increased expression of the β_2_-AR has beneficial effects under certain stress conditions. However, with *<u>lower level, chronic oxidative stresses</u>*, such as chronic DOX, pressure overload (14), dilated cardiomyopathy induced by MLP knockout (14), or diabetic cardiomyopathy (17), the β_2_-AR exerts cardiotoxic effects. In these settings, increased expression of β_2_-ARs exaggerates the deleterious phenotype, and β_2_-AR knockout mice are partially or fully rescued. We have shown that inhibition of β_2_-AR signaling during a chronic oxidative stress induces diverse signaling and metabolic compensations that combine to attenuate injury.

Thus, β_2_-AR signaling in cardiovascular disease may switch from cardioprotective to cardiotoxic depending not only on the type of stress but on the temporal nature of that stress. This temporal variability could hold for other cardiotoxic/cardioprotective signaling pathways, as few studies have compared cardiac stressors at both acute and chronic time points, and most animal models are extremely time-compressed approximations of human cardiac disease. Finally, β-blockers and β-agonists are amongst the most powerful and the commonly used drugs in cardiovascular medicine. Our findings of this β2-receptor signaling switch are highly relevant to the clinical use of subtype-specific β-antagonists and agonists, with their effects depending (more than is currently suspected) both on the underlying pathology and the chronicity of the stressor.

## Abbreviations

AMPK: AMP activated protein kinase
β-AR: β-adrenergic receptor
CaMKII: Ca^2+^/Calmodulin-dependent kinase II
DEG: Differentially expressed gene
DOX: Doxorubicin
FS: Fractional shortening
LV: Left ventricle
LVEDD: Left ventricular end-diastolic dimension
LVESD: Left ventricular end-systolic dimension
MLP: Muscle LIM Protein
ROS: Reactive oxygen species
TAC: Transverse aortic constriction

